# A Multi-omic Analysis of the Human Lung Reveals Distinct Cell Specific Aging and Senescence Molecular Programs

**DOI:** 10.1101/2023.04.19.536722

**Authors:** Ruben De Man, John E McDonough, Taylor S Adams, Edward P Manning, Greg Myers, Robin Vos, Laurens Ceulemans, Lieven Dupont, Bart M Vanaudenaerde, Wim A Wuyts, Ivan O Rosas, James S. Hagood, Namasivayam Ambalavanan, Laura Niklason, Kirk C Hansen, Xiting Yan, Naftali Kaminski

## Abstract

Age is a major risk factor for lung disease. To understand the mechanisms underlying this association, we characterized the changing cellular, genomic, transcriptional, and epigenetic landscape of lung aging using bulk and single-cell RNAseq (scRNAseq) data. Our analysis revealed age-associated gene networks that reflected hallmarks of aging, including mitochondrial dysfunction, inflammation, and cellular senescence. Cell type deconvolution revealed age-associated changes in the cellular composition of the lung: decreased alveolar epithelial cells and increased fibroblasts and endothelial cells. In the alveolar microenvironment, aging is characterized by decreased AT2B cells and reduced surfactant production, a finding that was validated by scRNAseq and IHC. We showed that a previously reported senescence signature, SenMayo, captures cells expressing canonical senescence markers. SenMayo signature also identified cell-type specific senescence-associated co-expression modules that have distinct molecular functions, including ECM regulation, cell signaling, and damage response pathways. Analysis of somatic mutations showed that burden was highest in lymphocytes and endothelial cells and was associated with high expression of senescence signature. Finally, aging and senescence gene expression modules were associated with differentially methylated regions, with inflammatory markers such as *IL1B, IL6R*, and *TNF* being significantly regulated with age. Our findings provide new insights into the mechanisms underlying lung aging and may have implications for the development of interventions to prevent or treat age-related lung diseases.

---

Age is a substantial risk factor in nearly all lung diseases^1^. Acute diseases such as pneumonia and ARDS, and chronic diseases such as bronchiectasis, chronic obstructive lung disease (COPD) and idiopathic pulmonary fibrosis (IPF) are both more common and more lethal in aged individuals^2^. While there has been significant progress in understanding the role of aging-related mechanisms in advanced lung disease^3-5^ and in describing the physiological effects of aging in the lung^6-8^, the cellular and molecular mechanisms that underlie the lung’s aging response remain poorly understood.

There has been limited knowledge generated concerning the cellular aging of the lung and much of it is in the context of disease^9^. Various studies have shown that pulmonary stem cell exhaustion and epithelial cell senescence are associated with advancing age and are implicated in the pathogenesis of IPF^3^. Mucociliary clearance and ciliary beat frequency were found to decrease with age and increase predisposition to pneumonia in the elderly^10^. Changes in ECM composition with age have also been described. Godin et al. reported increased collagen and decreased elastin and laminin in decellularized murine lung scaffolds^11^. More recently, Lee et al. reported age-associated fibrotic changes including increased density of collagen and decreased surfactant secretion^12^.

Studies of lung aging at single-cell resolution have been limited to date. Recent advances in sequencing technologies and the availability of larger datasets such as the Genotype-Tissue Expression (GTEx) Project^13^ have enabled the identification of novel markers of aging and senescence in other organ systems^14,15^. Chow et al. identified changes in bulk and single-cell RNAseq expression and relative cell type proportions with age in the context of SARS-CoV-2 susceptibility^16^. They observed increased transcriptional signatures associated with cell adhesion and stress responses. Angelidis et al. developed a mouse atlas of lung aging using single cell transcriptomics and mass spectrometry-based proteomics to determine the changes that occur with age^17^. To date, no comprehensive study of single-cell transcriptomic and epigenetic changes in the human lung with aging has been published.

In this work, we used an integrated multi-omic approach to determine cell type aging and senescence molecular programs in the human lung [Fig 1A]. We first identified gene co-expression networks in a large publicly available bulk RNA dataset^13^. Using single cell RNAseq data for validation, we showed that while some co-expression networks are broadly expressed, others reflected lung cell type-specific aging, with previously undescribed changes in alveolar cell subpopulations. Moreover, we defined an association between somatic mutation burden and expression of senescence-associated gene networks, connecting the damage accumulation hypothesis of aging with cellular senescence^18^. Finally, we identified epigenetic regulators of lung aging by assessing differentially methylated CpGs associated with co-expression networks.

**Figure 1:**
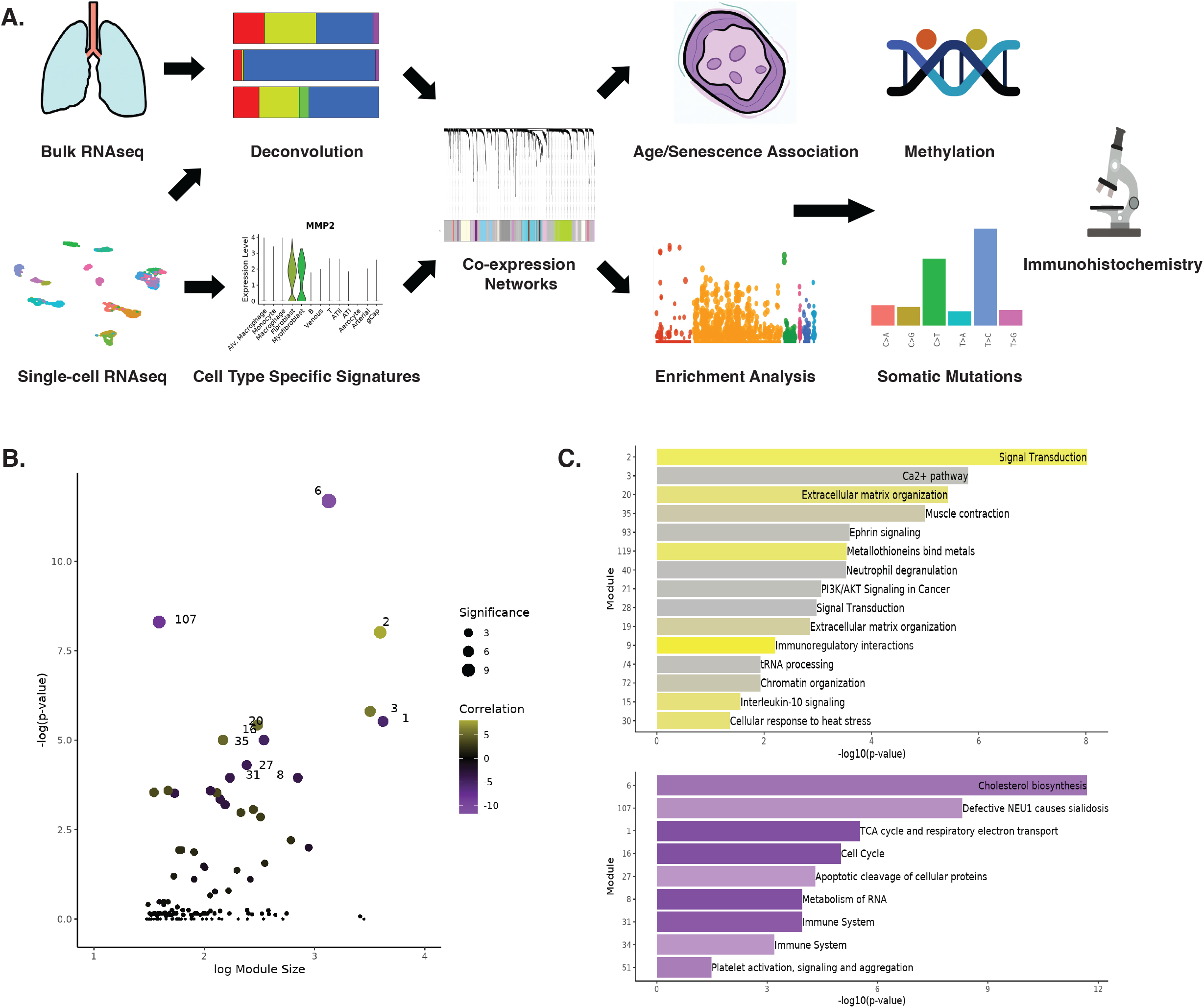
Study design and co-expression networks. (A) Gene co-expression networks were generated from GTEx bulk lung RNAseq data. Deconvolution of bulk RNAseq was performed using single-cell RNAseq data as a reference to determine cell type specificity of gene networks. Finally, enrichment for REACTOME terms and SenMayo signature was performed to characterize modules prior to downstream analysis. This schematic was generated with the assistance of DALL·E 2. (B) Co-expression networks developed in this study. The x-axis shows module size and the y-axis shows the FDR-corrected p-value for the correlation between the module eigengene and age. (C) Bar plots labelled with the top REACTOME term for age-associated modules. The top panel is positively correlated genes and the bottom panel is negatively correlated genes. The length of the bar represents the correlation of the module eigengene and age. The color of the bar corresponds to the hypergeometric test p-value for the REACTOME term enrichment.

## Results

### Gene co-expression network analysis reveals age-associated modules

To identify signatures of aging and senescence in transcriptomic data from lungs, we developed a gene co-expression approach that combines bulk and single-cell RNAseq data [Fig 1A]. Gene co-expression network analysis was performed using Weighted Correlation Network Analysis (WGCNA)^19^. Using bulk RNAseq dataset from the Genotype Tissue Expression Project, genes were grouped into 133 modules [Supp Fig 1] ranging from 30 to 4181 genes in size. Of these, 30 of 133 were significantly correlated with age (16 positive, 14 negative) [Fig 1B]. The molecular function of age-associated co-expression networks was assessed by enrichment of REACTOME pathways. Positively correlated modules were enriched for signal transduction, extracellular matrix (ECM) organization, and immunoregulatory interactions. Negatively correlated modules included TCA cycle and respiratory electron transport, cell cycle, and metabolism of RNA [Fig 1C]. This method uses gene co-expression analysis, which addresses some of the age-related increases in cell-to-cell variability and stochasticity ^20^. The results of enrichment analysis for co-expression networks were consistent with the those reported in in de Vries et al. ^21^ and Chow et al. ^16^, where linear models were applied to individual genes in the GTEx dataset to determine age differences in expression.

### Deconvolution of modules revealed that aging is associated with decreases in alveolar epithelial cells and increases in lung fibroblast and endothelial cells

To determine changes in cell-type-specific expression of aging modules, we performed deconvolution of bulk RNAseq data using our group’s previously published single-cell RNAseq dataset as a reference^22^ [Supp Fig 2]. Using the resulting cell-type proportion estimates for each sample, we sought to determine how the cell-type composition of the human lung changes with age. Cell-type proportions for each sample were correlated with age and Z-transformed [Fig 2A]. We observed significant decreases in the amount of both type I (Z=-0.28, FDR<0.001) and type II (Z=-0.30, FDR p<0.001) alveolar epithelial cells, consistent with previous published findings^16,23^. Endothelial cells, fibroblasts, and myofibroblasts all increased with age, suggesting that the cellular composition of the lung changes with advancing age. Specifically, aging appears to be associated with a reduced proportion of epithelial cells and an increase in endothelial and mesenchymal cells.

**Figure 2:**
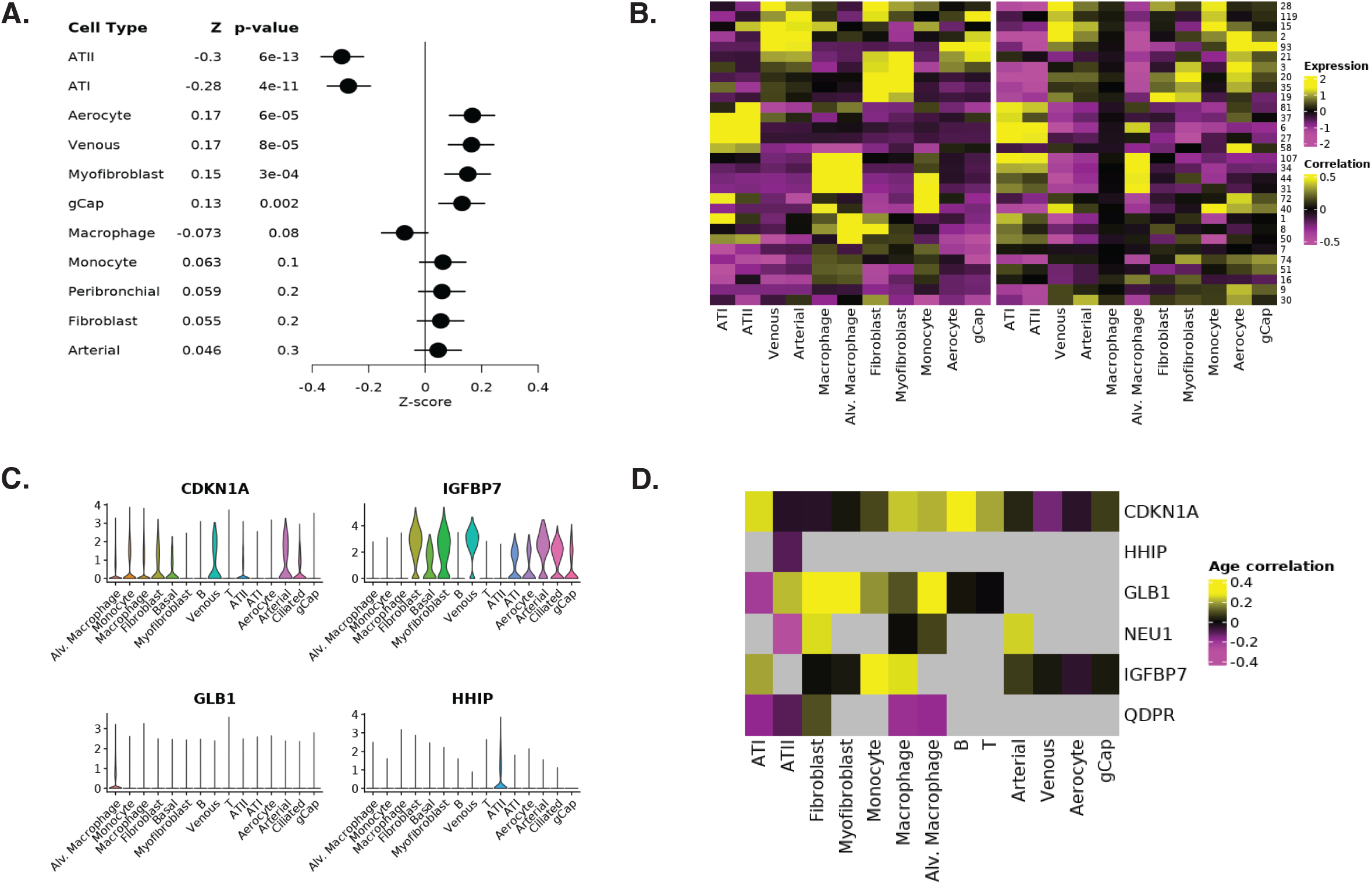
Age-associated gene co-expression modules correspond to specific cell types. (A) Changes in cell type proportions versus age. The Fisher Z-transformed correlation values for age versus cell type proportion and corresponding p-values are listed. (B) Cell-type specificity of 30 age-associated module eigengenes. The left panel shows the expression score for each module in each cell type in single-cell RNAseq data (38 samples). The right panel shows the correlation values between cell-type proportions and module eigengene in the deconvoluted bulk RNAseq dataset. (C) Violin plots for selected hub genes in age-associated modules. (D) Age correlation of selected hub genes in single-cell RNAseq data.

Next, we sought to determine whether co-expression modules corresponded to specific cell types. Cell type proportions for all samples were correlated with the module eigengene for each module, revealing patterns of cell type specificity. To assess whether this cell-type deconvolution was successful, we calculated expression of each module in our single-cell RNAseq dataset. Heatmaps of both cell type associations based on deconvolution and single-cell expression scores were plotted side by side, demonstrating the accuracy of this approach [Fig 2B]. Notably, agreement between both methods was present for the majority of cell-types, with the exception of cells with low representation in our bulk RNAseq dataset (macrophages, B/T cells, peribronchial cells, and ciliated cells) [Supp Fig 2].

To identify specific genes that drive module function, hub genes were identified for each of the age and senescence-associated modules. Hub genes were defined as those that had a high module membership (correlation of gene expression to ME) and high correlation with age based on expression in GTEx RNAseq data. Notable hub genes included *IGFBP7* in the ECM module, *QDPR* in the mitochondrial module, and *HHIP* in the cholesterol biosynthesis module [Fig 2C]. *GLB1* which encodes SA-B-Gal, one of the gold standard markers for cellular senescence^24^, and *NEU1*, which encodes a lysosomal sialidase implicated age-related neurodegeneration^25^, were hub genes for the lysosome module. To confirm the association of these hub genes with aging, age correlation was calculated directly in our single-cell RNAseq dataset [Fig 2D]. *IGFBP7* was expressed in most cell types and positively correlated with age in AT1 and immune cells (FDR p<0.001). *GLB1* and *NEU1* were expressed and positively correlated with age in mesenchymal and immune cells (FDR p<0.001). *QDPR* was negatively correlated with age in epithelial cells and macrophages (FDR p<0.001). Finally, *HHIP* was negatively correlated with age and specific to AT2 cells (r=-0.11, FDR p<0.001). Hence, we used single-cell RNAseq data to confirm the cell-type specificity and age association of module hub genes.

### Alveolar microenvironment aging is characterized by decreased AT2B cells and reduced surfactant production

The cholesterol biosynthesis module was the most strongly negatively (r=-0.31, FDR p=2e-12) correlated with increased age [Fig 1B]. This module contained *HHIP* as a hub gene, a gene that has been implicated in the pathogenesis of COPD and is expressed at lower levels in COPD subjects based on human single-cell RNAseq^26^. This module and its hub gene were primarily expressed in AT2 cells [Fig 3A, 3B]. We examined the expression of *HHIP* in both bulk and single-cell RNAseq datasets and confirmed that its expression is negatively correlated with age (r=-0.34, p=9e-17) [Fig 2D, 3C]. Finally, other established AT2 cell markers (*SFTPA1, SFTPA2, SFTPC*) were found to correlate with *HHIP* expression (r =0.30 to 0.37, FDR p<0.001) and were negatively correlated with age (−0.19 to -0.09, FDR p<0.001) [Fig 3D]. Taken together, these findings suggest that *HHIP* is a central gene in a molecular program that leads to reduced surfactant production in AT2 cells. When examining single-cell data for AT2 cells independently, we observed two subpopulations that formed independent clusters on a UMAP plot [Fig 3E], similar to what has been reported previously by our group and others^26-28^. Travaglini et al. described two subtypes: bulk AT2 (AT2B) cells that have high expression of surfactants and correspond with a functional phenotype, and signaling AT2 (AT2S) cells that have lower levels of surfactant and correspond with a more stem-like progenitor phenotype ^27^. In our dataset, we confirmed that AT2S cells exhibited reduced expression of surfactants and the surfactant-associated gene module compared to AT2B cells [Fig 3E]. Additionally, AT2B cells were associated with lower subject age (41 versus 49, FDR p<0.05) and exhibited higher expression of *HHIP* (0.44 versus 0.14, FDR<0.05) [Fig 3E,3F]. Hence, the proportion of *HHIP*-expressing, surfactant-producing AT2B cells decreased with age. To confirm the association between *HHIP* expression and age, we performed immunohistochemistry in aged and young FFPE sections. HHIP was stained with surfactant protein C (SPC), a marker for type II alveolar epithelial cells. We observed co-localization of HHIP and SPC, confirming that HHIP is primarily expressed in AT2 cells [Fig 3G]. We further sought to validate the negative correlation between HHIP expression and age that we observed at the transcriptomic level. We quantified the proportion of SPC-positive cells that also stained positive for HHIP. We observed a significant difference between young and aged subjects (0.55 versus 0.31, t=2.95, p=0.016) [Fig 3H]. Taken together, these results indicate that AT2 cells not only decrease in number with age and exhibit reduced expression of gene networks central to surfactant production and normal progenitor function, but also that the subpopulation functionally involved in maintenance of the alveolus declines. These findings potentially explain the lung’s increased predisposition to injury with age.

**Figure 3:**
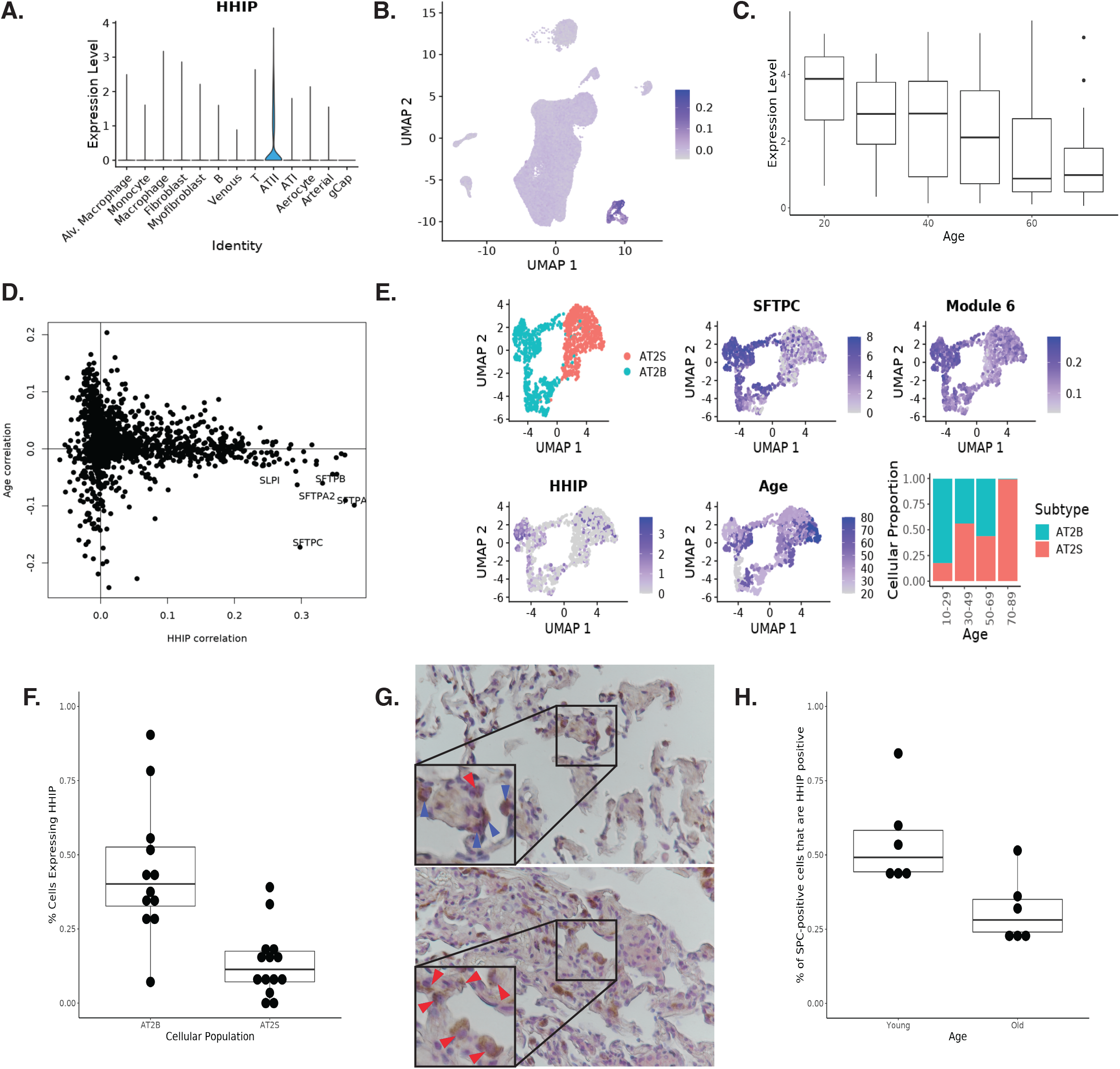
HHIP expression is reduced in ageing and correlates with markers of ATII cells. (A) Violin plot of HHIP expression across cell types. (B) UMAP representation of HHIP expression. (C) HHIP expression declines with age (r=-0.34, p=9e-17). The x-axis is age and the y-axis is average expression of HHIP in bulk RNAseq. (D) ATII cell markers correlate with HHIP and are negatively correlated with age in single-cell RNAseq data. (E) UMAP representation of AT2 cells, revealing AT2A and AT2B subpopulations. (F) Proportion of cells in each sample expressing HHIP in AT2A vs AT2B subpopulations based on scRNAseq data. (G) Representative immunohistochemistry images for dual SPC/HHIP staining. The top panel is a young subject and the bottom panel is an aged subject. SPC was stained with DAB (brown), HHIP was stained with AP Red (pink). Red arrowheads indicate SPC positive cells and blue arrowheads indicate dual positive, co-stained cells (H) The proportion of dual positive cells among SPC positive cells in young versus aged subjects.

### SenMayo captures cells expressing a senescence gene signature but does not correlate with age

Next, we were interested in determining whether we could identify senescence-associated modules by using previously identified senescence gene signatures. Modules were assessed for enrichment of six established senescence gene signatures^29,30^. Fisher’s Exact testing was performed to compare enrichment of all six gene lists across all modules. Strong overlap was observed across known senescent gene lists [Fig 4D, Supp Fig 3]. Interestingly, module 15 was the only module that was significantly enriched (p<0.05) for all six senescence lists. Because the SenMayo gene signature outperformed all other gene lists in its ability to detect senescent cells^14^, we evaluated SenMayo as a potential gene signature for further analysis of co-expression modules. A SenMayo score was calculated for all cell types in a single-cell RNAseq dataset to determine whether this gene signature can accurately identify cells with senescent phenotype [Fig 4A]. High SenMayo cells were identified using an arbitrary SenMayo score threshold. These cells were then plotted using a UMAP representation, showing that they cluster together [Fig 4B]. To confirm that SenMayo score was characterizing senescent cells and not cells with an inflammatory phenotype, we also examined expression of canonical senescence markers in this dataset. Expression of *CDKN1A*, a known marker of senescence not included in the SenMayo list, was found to be increased in high SenMayo scoring cells [Fig 4C]. This subpopulation consisted primarily of fibroblasts, macrophages, and endothelial cells [Fig 4E]. Notably, SenMayo score was not significantly correlated with age in any cell type (FDR p>0.05).

**Figure 4:**
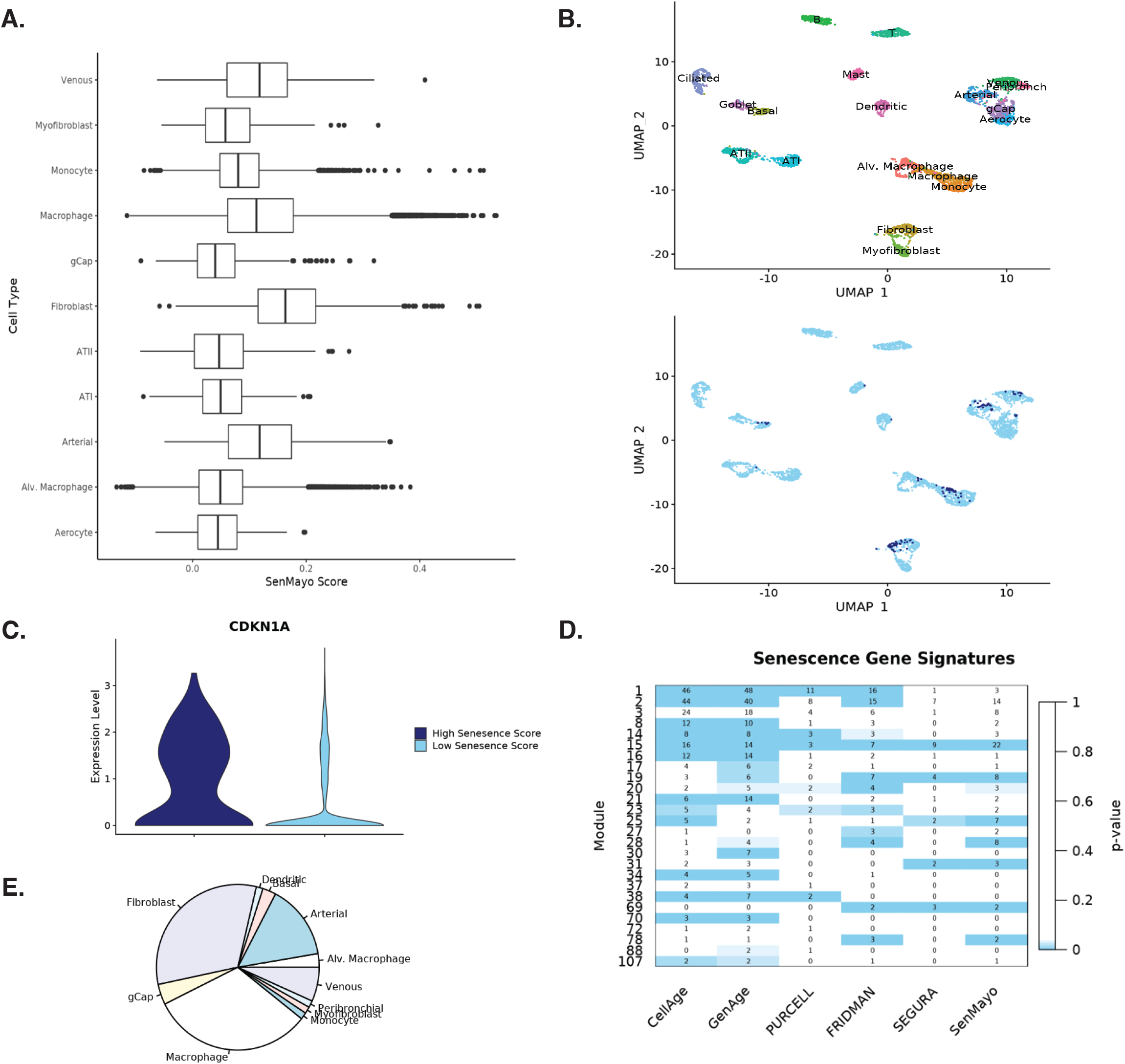
SenMayo scoring can be used to identify senescence signature in RNAseq data. (A) SenMayo score for various cell types in single-cell RNAseq data. (B) High SenMayo cells cluster together on a UMAP plot. The dataset was downsampled to include up to 250 cells per cell type. The top panel depicts cells clustered by cell type. The bottom panel shows high SenMayo cells, highlighted in dark blue, clustering together. (C) High SenMayo cells have higher expression of independent senescent markers such as CDKN1A. (D) Breakdown of cell type composition of high SenMayo cells. (E) Heatmap showing module enrichment for senescent signature for 6 senescence gene lists. Heatmap intensity corresponds to the FET p-value.

### Senmayo identifies senescence-associated co-expression modules that have distinct molecular functions

After establishing that SenMayo could be used to identify senescent features in our datasets, we used this signature to determine which modules were associated with cellular senescence. Of the 30 modules correlated with age, six modules were also enriched for SenMayo gene signature (FET p<0.05) accounting for 59 of the 125 SenMayo genes [Fig 5A]. These modules had distinct functional enrichment for REACTOME terms [Fig 5B]. Additionally, they exhibited cell-type-specific expression, both in single-cell data and based on deconvoluted proportions [Fig 5C, 5D]. The module that included the greatest number of SenMayo genes (23 genes including *CXCL1/CXCL2/CXCL3, IL1A/B, IL6, TNF, SERPINE1*, and *VEGFA*) also included p21/*CDKN1A*, a hallmark of senescence, as a hub gene. As we previously noted, this was the only module significantly enriched for all six established senescence lists. Deconvolution correlated this module (15) with vascular and monocytic cells, with enrichment for cytokine/interleukin signaling pathways. p21/*CDKN1A* was also significantly correlated with age in these cell types in the single-cell RNAseq dataset (p<0.05). Two modules (19, 20) correlated with fibroblast and myofibroblast cell types and were enriched for ECM pathways. Genes included *IGF1* (19), IGF binding-proteins, and matrix metalloproteinases. Another two modules (28, 31) were enriched in immune pathways and corresponded to monocytes/endothelial cells and alveolar macrophages, respectively. Module 31 was negatively correlated with age and its top hub gene, *SYK*, encodes a signaling molecule associated with immune deficiency and dysregulation. Finally, the module most strongly correlated with age (2, r=0.26, p<0.05) contained Wnt signaling genes and p16/*CDKN2A* as a hub gene. This module demonstrated enrichment for signal transduction pathways and chromatin modifying enzymes/histone deacetylases and was associated with endothelial cells. Interestingly, other hub genes for this module were associated with DNA damage response and oncogene-induced senescence. The module also contained genes such as *ATM* (a gene which is activated by DNA double strand breaks), *MSH2*, as well as known oncogenes such as *KRAS* and *RAF1*^*31*^.

**Figure 5:**
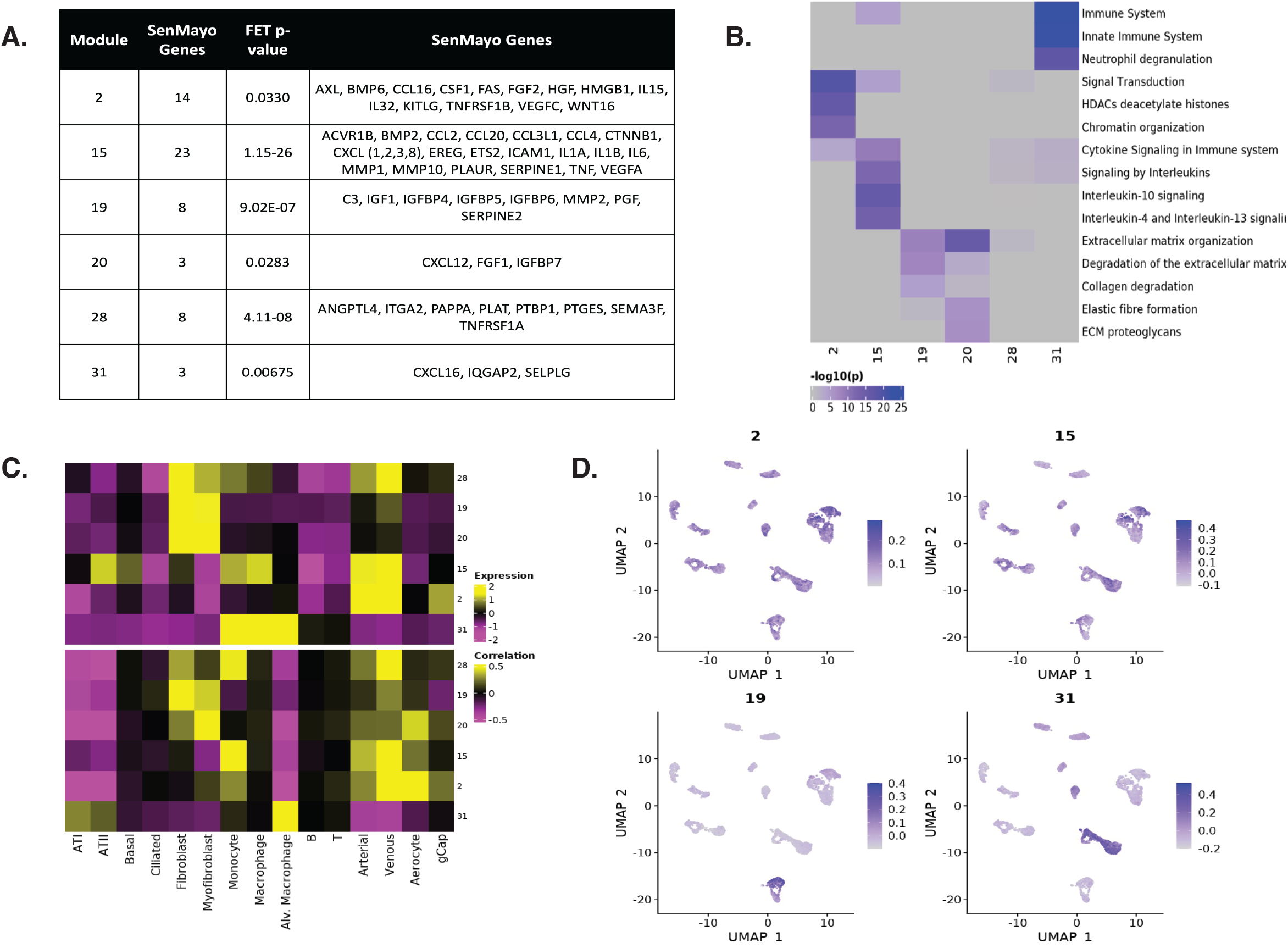
Deconvolution reveals cell-type specific signatures of senescence. (A) Table of senescence-associated modules. This table includes the number and identities of SenMayo genes present in each module. (B) Heatmap showing enrichment of top 3 REACTOME terms for each module. (C) Cell-type specificity of 6 senescence-associated module eigengenes. The top panel shows the expression score for each module in each cell type in single-cell RNAseq data. The bottom panel shows the correlation values between cell-type proportions and module eigengene in the deconvoluted bulk RNAseq dataset. (D) Cell-type specificity of senescence-associated modules on UMAP.

To confirm that modules were biologically relevant and generalizable to other datasets, module preservation analysis was performed. Module Z-summary scores (see methods) were calculated using data from the Lung Tissue Research Consortium (LTRC) as a validation dataset. Using the threshold established in [Supp Fig 1], we observed that 5 out of 6 senescence modules were conserved in this second dataset, indicating replication and biological relevance. We noted that the module that did not meet the threshold (20) was enriched for similar REACTOME terms and contained related genes to module 19, indicating that this was module may have been segregated due to the relatively sensitive module detection parameter (see methods).

### Somatic mutation burden is concentrated in lymphocytes and endothelial cells and is associated with high expression of senescence signature genes

To further understand the relationship between aging and cellular senescence, we analyzed somatic mutation data from a previously published GTEx study^32^ in order to determine the role of somatic mutation accumulation in the expression of senescence gene modules. Somatic mutation data was available for 321 of the original 572 subjects which we integrated with the gene module expression data [Fig 6A]. Two senescence modules (2,15) were among the most correlated with somatic mutation burden (133 total modules) [Fig 6B, 6C]. Interestingly, these were the two modules contained *CDKN1A* and *CDKN2A*, supporting the association of DNA damage with senescence. Among senescence-associated gene modules, module 2 had the strongest correlation with mutation burden and with age [Fig 6B]. In addition to *CDKN2A*/p16, this module also contained DNA damage response associated genes, including *ATM, ATR, HMGB1*, and *MSH2*. To determine whether specific cell types were prone to somatic mutation accumulation, deconvoluted cell type proportions for each sample were correlated with global somatic mutation burden. We found arterial/venous endothelial cells and T cells were significantly associated with mutation burden (FDR p<0.05), indicating that these cell types had increased accumulation of somatic mutations in this dataset [Fig 6D]. We hypothesized that cell types with greater association with somatic mutation burden would also express DNA damage response genes at higher levels in aged cells. To confirm this, and to validate the association of DNA damage response genes in this module with age, we calculated correlation with age in the single-cell RNAseq dataset [Fig 6E]. We observed higher correlation of markers such as *ATM* and *ATR* with age in endothelial cells (FDR p<0.001) and lymphocytes (FDR p<0.001), consistent with the higher mutation burden in these cell types. This module was also expressed primarily in endothelial cells [Fig 5C]. Taken together, these results suggest that there are specific gene networks that link DNA damage accumulation with cellular senescence. Additionally, specific cell types, including endothelial cells, appear to be more prone to damage accumulation.

**Figure 6:**
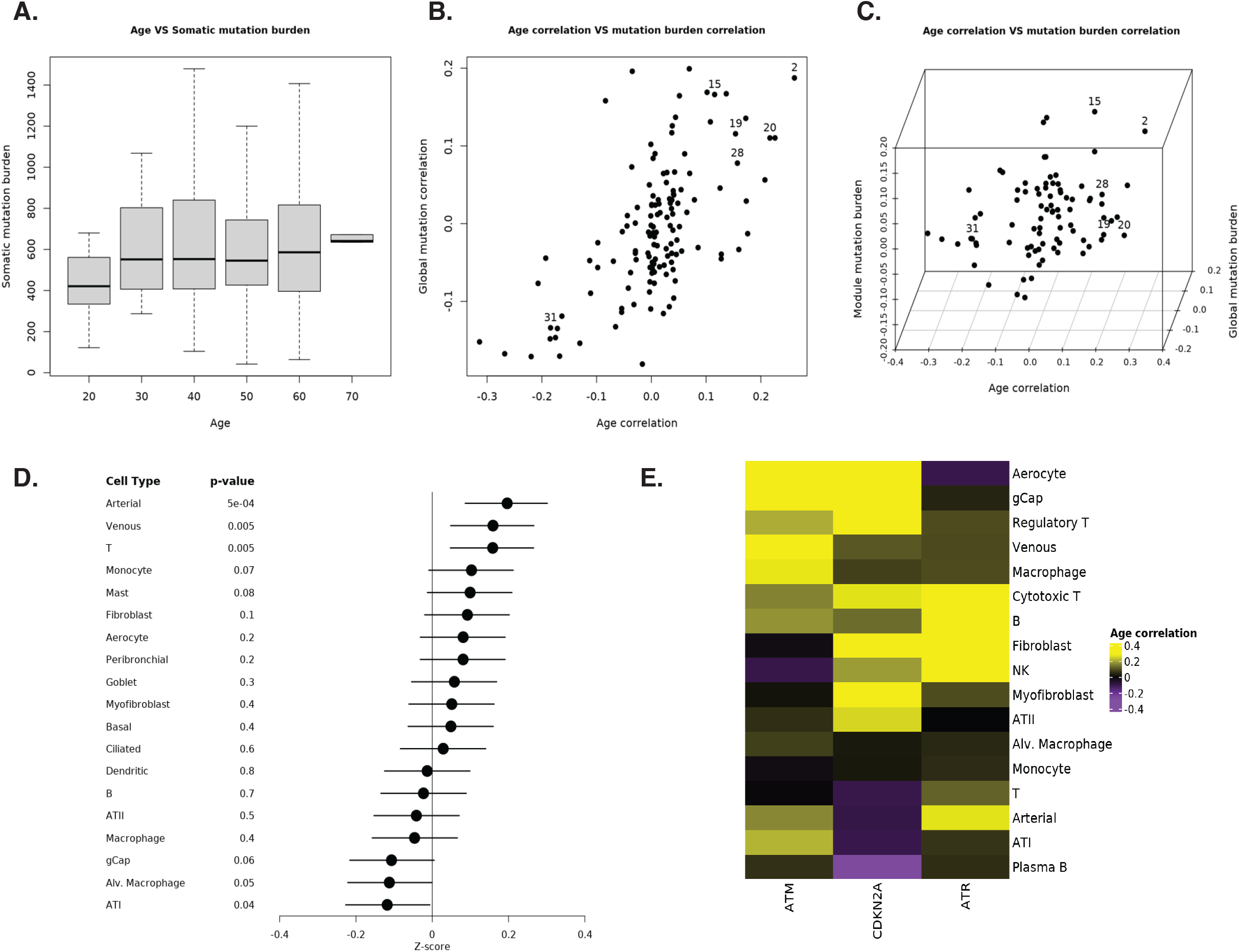
Senescence signatures are associated with somatic mutation burden and DNA damage response. (A) Somatic mutation burden increases with age. (B) Plot of age correlation versus global mutation burden. The x-axis shows correlation values between module expression and age correlation. The y-axis shows correlation values between module expression and global mutation burden. (C) 3D plot of age correlation versus global mutation burden versus module mutation burden. (D) The Fisher Z-transformed correlation values for mutation burden versus cell type proportion and corresponding p-values. (E) Age correlation of DNA damage response genes in single-cell RNAseq data.

### Aging and senescence gene expression modules are associated with differentially methylated regions

To determine whether epigenetic modifications play a role in module control, we analyzed whole methylome data from 27 samples with ages ranging from 29 to 79. Methylation data corresponding to 865918 CpGs were available. We integrated CpGs with their corresponding gene annotations and restricted analysis to CpGs that were linked to genes that were present in one of age (n=347956) or senescence-associated gene modules (n=111351) [Fig 7A, 7B]. Only CpGs that had β values for all subjects were retained. Approximately an equal number of CpGs were positively and negatively correlated with age [Fig 7C]. 10 senescence CpGs were significantly correlated with age (FDR p-value <0.05). Among these, the top two were both associated with *ROCK2* [Fig. 7D], a gene that has been implicated in cardiac fibrosis and age-related aortic stiffening ^33^. An additional two markers corresponded to *ITGA1*, a key regulator of Cell-ECM communication and the resulting dynamic reciprocity^34^. To better distinguish genes whose expression is strongly under epigenetic control, we identified genes with a significant proportion of CpGs showing strong association with age. Fisher’s Exact Test was performed to find genes that were enriched with significantly age-correlated CpGs. Given the role of module 15 in controlling induction of senescence-associated secretory phenotype (SASP), we applied this analysis to genes in this module. Expression of 207 genes was positively correlated with age and 58 genes were negatively correlated. Genes that were positively correlated with age had a higher proportion of enrichment for hypomethylated CpGs (p<0.05). Interestingly, the most enriched genes included well-known immune signaling genes such as *IL1B, IL6R*, and *TNF* [Fig 7E]. Other genes included *ETS2*, a gene involved in senescence signaling downstream of p38MAPK^35^. Taken together these finding support a strong association of methylation epigenetic changes with age related changes in inflammatory gene expression changes.

**Figure 7:**
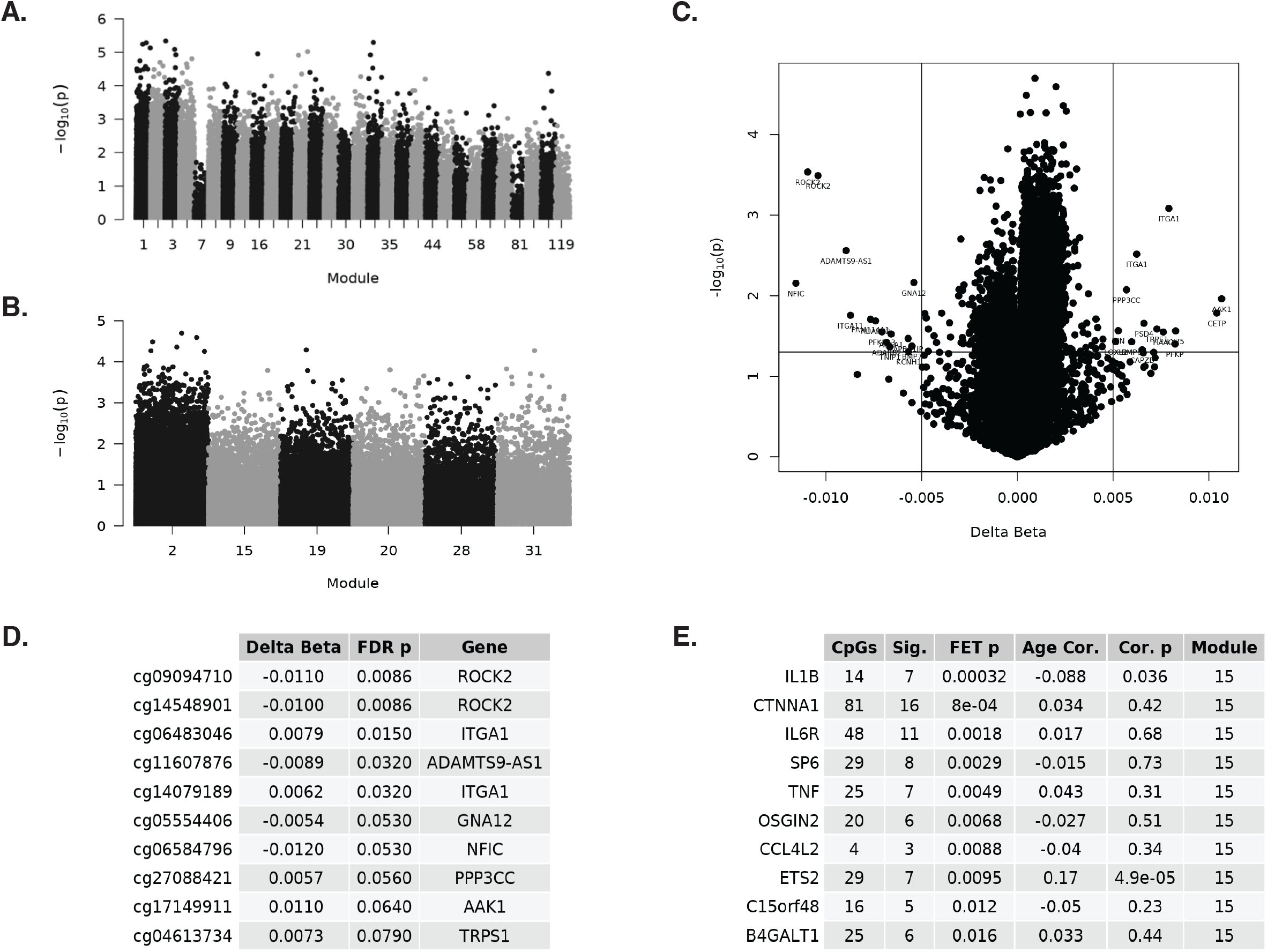
Senescent genes are regulated epigenetically. (A) Manhattan plot of all CpGs with a gene annotation in age-associated modules. (B) Manhattan plot of all CpGs with a gene annotation in senescence-associated modules. (C) Plot of delta beta values and log transformed p-values for all senescence-associated CpGs. (D) The top ten senescence-associated CpGs, sorted by FDR p-value for age correlation. (E) The top ten genes in the SASP module (15), sorted by FET p-value for enrichment of age-associated CpGs.

## Discussion

In this study, we identified aging and senescent signatures in the aged human lung by performing integrated analysis of various large datasets. We developed co-expression networks using bulk RNAseq data from the GTEx project and validated this using several independent datasets: a single-cell RNAseq dataset, bulk RNAseq dataset from the LTRC, and FFPE lung tissue sections also from the LTRC. Gene modules reflected hallmarks of aging, including mitochondrial dysfunction, inflammation, and cellular senescence. Deconvolution revealed significant shifts in the cellular composition of the aged lung, with a decrease in epithelial cells and increased fibroblasts and endothelial cells. The module most strongly correlated with age (r=-0.31, FDR p=2e-12) was related to surfactant production and was expressed in AT2B cells, which declined in proportion with age. We examined gene networks enriched for SenMayo signature, and found that these were associated with specific cell types and had distinct molecular functions, including cell signaling, damage response pathways, SASP, and chromatin regulation. Analysis of somatic mutations revealed an association between elevated mutation burden and expression of senescence gene networks. Finally, these aging and senescence gene expression modules were associated with differentially methylated corresponding to inflammatory markers and genes such as *ROCK2* and *ITGA1*. Together, the results of this multiomic analysis revealed cell specific changes associated with lung aging and their dependence and independence of cellular senescence.

Our integrated analysis allowed us to obtain novel observations regarding the cellular changes that occur in the alveolar microenvironment with age. Our deconvolution of bulk lung RNAseq data indicated that the proportion of type I and type II alveolar epithelial cells decreases with age. Previous studies have suggested a decrease in AT2 cells with age, particularly in the context of severe COVID-19^16^ and pulmonary fibrosis^12^. Using gene co-expression analysis, we identified a module associated with surfactant production in type II alveolar cells that was negatively associated with age. *HHIP*, one of the genes most strongly associated with susceptibility to COPD in human GWAS studies and functionally with maintaining normal lung function and alveolar structures in mice ^*36-38*^, was a hub gene for this module. This finding suggested an age-related change in AT2 cell subpopulations because *HHIP* is primarily expressed in one subpopulation identified as AT2B cells^26,27^. When examining single-cell data, AT2 cells were comprised of two major groups: AT2B cells that have high expression of surfactants and correspond with a functional alveolar maintaining phenotype, and AT2S cells that have lower levels of surfactant and correspond with a more stem-like progenitor phenotype^28^. Our scRNAseq data, as well as our immunohistochemistry validation confirmed decreased expression of *HHIP* with age, as well as a decrease in AT2B cells in the aged lung. Interestingly, among AT2S cells, surfactant gene expression was also decreased. Thus, our analyses uncover a both a specific decline in AT2B with age, as well as a general decline in surfactant gene networks among alveolar epithelial cells. While the specific role of *HHIP* in this context is unknown, the decline in surfactant transcriptional programs and the relative decline in AT2B may have important implications in explaining the increased predisposition of the aged lung to alveolar injury and diseases such as ARDS and pneumonia^9,39^.

The co-expression networks in this study identified age-associated molecular programs which reflected some of the hallmarks of aging^40^. Positively correlated modules were enriched for terms such as signal transduction, ECM organization, and immunoregulatory interactions, while terms for negatively correlated modules included TCA cycle and respiratory electron transport, cell cycle, and metabolism of RNA. These findings are consistent with other studies that examined lung aging signatures in the GTEx dataset on a bulk level. de Vries et al.^21^ examined whole genome mRNA on lung tissue samples and identified 3509 genes that changed with age. This gene expression signature was validated by determining significance in the Genotype-Tissue Expression (GTEx) dataset^21^. Another study examined gene expression changes in the GTEx project identified 876 genes that changed with age. Genes associated with lung aging were enriched for mitochondrial and lysosomal pathways^41^. Hence, our co-expression networks confirmed known changes associated with lung aging, while identifying a new pattern of alveolar epithelial cell aging.

After assessing the changes in the transcriptome with age, we analyzed modules that were also associated with senescence. The top senescence-associated module included *CDKN1A* as a hub gene and was closely co-expressed with interferon signaling genes *STAT3, IL4R, IL1R1, CCL2*, and *IL6*, suggesting its central role in connecting cell cycle and SASP production. Another module (2) was expressed in most cell types and contained *CDKN2A*. Enrichment analysis showed that this module contained a wide variety of genes involved in DNA repair and DNA damage response. This supports the idea of specific gene networks that link cellular damage response and cellular senescence^18^. Notably, *CDKN1A* and *CDKN2A* were associated with different modules in this analysis, indicating their likely contributions to different senescence signaling pathways. This has been previously reported in the literature^42^. Our findings suggest that p16 and p21 have different roles in the induction and maintenance of cellular senescence in the lung. Whereas p21 appears to be more directly related to the SASP, p16 is linked with DNA damage response and oncogene induction. Additionally, we identified a senescence module specifically expressed in fibroblasts, which included known senescence gene *IGF1* and many of its associated binding proteins. This module also contained many collagen genes and matrix metalloproteinases, suggesting a link between IGF signaling, lung fibroblast function and dysregulation of extracellular matrix deposition associated with aging and senescence, consistent with recent findings ^43^, but these findings will require additional validation. Interestingly, while age-associated modules were enriched for SenMayo signature, SenMayo score was not significantly correlated with age in any cell type. Previous studies have shown significant heterogeneity in the relationship between senescence and chronological age across different tissues^44,45^. Moreover, the magnitude of the relationship depends on the marker used to detect senescence. Notably, the association between age and senescence has not been adequately studied in terminally differentiated cells, such as the AT1 cells that largely comprise the alveolar epithelium^44,46^. The lack of evidence for such an association in this study suggests that effects of cellular effects of lung aging distinct from the effects of cellular senescence in certain lung resident cell subpopulations. The differential somatic mutation burden and epigenetic changes as discussed below, suggest that this is indeed is the case, but more detailed studies are required.

Previous studies have suggested that cellular senescence is associated with macromolecular damage accumulation, with processes such as telomere shortening, oxidative stress, and somatic mutations contributing^18,47^. We identified senescence signatures in the aged lung and observed a strong association between somatic mutation burden and expression of senescence-associated genes in our samples. We determined that somatic mutations rates are highest in endothelial cells, which also had the highest expression of senescence signature. Notably, endothelial cell senescence has been associated with age-related lung diseases such as COPD^48^. Similar analysis has been performed on single-cell human pancreas data, showing a correlation between mutation load and *CDKN2A* expression in endocrine cells^49^. However, this has not been reported in the lung. Together, these findings provide support to the long-standing hypothesis that accumulation of macromolecule damage, including DNA damage, leads to cellular senescence in the aged human lung^18^. Moreover, we show the previously undescribed observation that specific cells, especially endothelial cells, within the aged lung are more susceptible to mutation accumulation and cellular senescence.

Finally, epigenetic regulation of aging and senescence signatures was examined by identifying variably methylated CpGs and integrating these with gene expression levels. We observed several CpGs that were differentially methylated with age. Many of these CpGs corresponded to senescence gene co-expression modules, suggesting that regulation of cellular senescence is at least in part by epigenetic mechanisms. This has been shown in a variety of cancer studies^50^. The top two senescence-associated CpGs that correlated with age both had *ROCK2* as an annotation, which is involved cardiac fibrosis and age-related aortic stiffening ^33^. Additionally, we saw evidence of epigenetic regulation of SASP, with genes such as *IL1B, IL6, TNF*, and *ETS2* having significant enrichment of age-correlated CpGs. These epigenetic regulators were associated with modules primarily expressed in endothelial cells and immune cells. Hence, we identified novel epigenetic regulators of cellular senescence showed that they corresponded to specific lung cell types.

The study has some limitations. There are limitations to bulk RNA sequencing deconvolution; Some cell types were less represented in our deconvoluted bulk RNAseq dataset [Supp Fig 2], also deconvolution from bulk data may represent decreased activity rather than a change in composition of cell types. These limitations may make the cell inferences inaccurate. We addressed this by using our single single-cell RNAseq data showing that examining module expression directly in single-cell RNAseq data recapitulates the cell-type specificity observed with deconvolution, and identifying cell subpopulation specific changes and validating them as we did in the case of the decline in AT2B with aging. Our method of determining senescence-associated gene co-expression networks relied on previously published senescent gene lists, which likely do not fully reflect the cell-type specific mechanisms of cellular senescence in the aged human lung. However, we confirmed that SenMayo identifies senescent signature by assessing the expression of independent markers of senescence, such as *CDKN1A*.

In summary, our study represents the most comprehensive multi-omic study of lung aging to date; we have identified novel cellular and molecular programs associated with aging and cellular senescence in specific cell types and discovered distinct changes in cellular composition, somatic mutation burden and epigenetic changes in the aged human lung. Further studies will be needed to further characterize and describe the temporal and spatial dynamics of these changes and define the role of these changes in the enhance disease predisposition and reduced resilience.

## Methods

### Bulk RNAseq Datasets

Lung-specific bulk RNAseq data was downloaded from the Genotype-Tissue Expression (GTEx) Portal (https://gtexportal.org/home/datasets). GTEx consists of samples from 54 non-diseased tissue sites across nearly 1000 individuals. Tissue was collected from postmortem/organ procurement cases. The data consisted of 572 samples with an age range of 20 to 79 years. Because the age of GTEx subjects was reported in 10-year ranges, the mean value of these ranges was used for subsequent analysis. The normalized gene expressions were log2 transformed.

A second bulk lung RNAseq dataset from the Lung Tissue Research Consortium (LTRC) was processed in parallel with the original dataset and used to assess for module preservation across datasets. This dataset was downloaded from NCBI GEO GSE47460 (https://www.ncbi.nlm.nih.gov/geo/query/acc.cgi?acc=GSE47460). The same parameters for data pre-processing were applied to this dataset. This data set consisted of 91 samples, with an age range from 32 to 87 years.

### Weighted gene co-expression network analysis

WGCNA was conducted using R/Bioconductor to identify modules in the normalized bulk RNAseq dataset. Analysis parameters were adjusted: sign of correlations between neighbours (TOMtype and networkType=‘signed’), and module detection parameter (deepSplit=2). Modules were identified by number in order of decreasing module size. Module eigengene (ME) was calculated as the first principal component of gene expression for the module. Module association with age was calculated using the module eigengenes (first principal component of expression profile). Pearson correlation adjusted for multiple comparisons by FDR. Module membership, a measure of the association of a gene to its module, was determined by Pearson correlation of gene expression to ME and used to rank module connectivity.

Module preservation analysis was performed to determine whether modules defined in the GTEx dataset persisted in the LTRC dataset. This was done using the “modulePreservation” function in WGCNA, which calculates module connectivity preservation statistics including correlation of correlations and correlations of eigengene-based connectivity kME. Significance of each module preservation measure significance was calculated using the observed value and the permutation Z score.

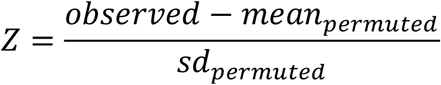

Z scores are a measure of module preservation compared to a random sample of genes. A composite measure called Z-summary was used to determine preservation. Modules were considered preserved if their Z-summary score was greater than 10, indicating strong evidence of preservation.

### Enrichment Analysis & Hub Genes

Enrichment of REACTOME pathways was determined using GProfiler in R. Modules were then assessed for the presence of genes from the SenMayo gene list^14^ using Fisher’s Exact Test (FET).

Hub genes were defined as those that had a high module membership (correlation of gene expression to ME) and high correlation with age based on expression in GTEx RNAseq data. In order to validate these hub genes, the correlation of hub gene expression with age was calculated in the single-cell RNAseq dataset. Expression values were calculated for each cell-type individually, and only cells with a minimum expression value were included in analysis.

### Single-cell RNAseq

Single cell RNAseq data reported in our group’s previously reported IPF Cell Atlas^22^ were used for this study. Healthy controls consisted of 38 samples from 28 subjects. This dataset consisted of a total of 96083 cells, with an age range from 20-80 years. scRNAseq data analysis was performed using the standard Seurat pipeline. Gene expression values were normalized by “NormalizeData” method. The top 2,000 variable genes were identified by the “FindVariableFeatures” function. After scaling gene expressions, a linear dimensional reduction was performed using the variable genes by “RunPCA” functions, which generated 30 principle components. Clustering was performed using the “FindNeighbors” function, which uses a K-nearest neighbor (KNN) graph. The “FindClusters” function was subsequently applied to optimize modularity by the Louvain algorithm. The resolution parameter for the clustering granularity was set to 0.05. The “RunUMAP” function was applied for nonlinear dimensional reduction and cluster visualization.

The same parameters were used to cluster AT2 cells independently, with the exception of the resolution parameter, which was reduced to 0.01. A total of 1251 AT2 cells were present in the dataset.

Module and gene age correlation and cell-type specificity were validated by calculating expression in the single-cell dataset (expression of each gene list subtracted by expression of random control feature sets). Module expression and SenMayo score were calculated using the “AddModuleScore” function in Seurat, which averages the expression of each gene list subtracted by expression of random control feature sets. Correlation between SenMayo score and age was calculated by determining the average SenMayo score per cell type, per subject, and correlating with subject age.

To identify marker genes of each cell type, the “FindAllMarkers” function from Seurat was applied to identify differentially expressed genes using a Wilcoxon Rank Sum test. Only significantly upregulated genes (FDR < 0.05) with 0.25 log fold change and 0.25 minimum expression fraction were retained as marker genes.

### Deconvolution

The GTEx bulk RNAseq dataset was deconvoluted to determine cell type proportions using MuSiC in R^51^. MuSiC uses support vector regression on gene expression profiles using reference gene expression signatures. Our single-cell RNAseq dataset was used for this purpose. The cell type composition was determined for each sample from the signatures in the original expression profiles. The resulting cell type proportions were correlated with the module eigengenes.

### Somatic Mutations

Somatic mutation data was acquired from Garcia et al.^32^. Mutations in this study were calculated by mapping raw RNAseq reads to the reference genome Hg19 and deploying a comprehensive mutation calling pipeline. False positive mutation calls were minimized by accounting for sequencing errors, RNA editing events, germline variants, and other sources of error. Somatic mutation burden for the present study was calculated using the sum of somatic mutation counts for each sample^32^.

### Methylation

Donor lungs that were not suitable for transplantation were collected at University Hospital Leuven, Belgium, and used as “healthy” controls. 28 donor lungs were used ranging in age from 20-80, with 7 females and 21 males. This study was performed with approval from the hospital ethical committee (S51577). Donor lungs were obtained as previously described ^52,53^. Whole methylome data was performed on all 28 samples. Methylation was assessed using the Infinium Human Methylation 450K Bead Chip (Illumina, Inc). A total of 865918 CpGs were present in the dataset. At each CpG site, methylation was reported as a β value, which is the proportion of signal obtained from the methylated beads over the sum of signal from all beads. β values ranged from 0 (no methylation) to 1 (full methylation). Pre-processing was performed per the manufacturer’s protocol. Data were normalized to internal controls. Signals corresponding to probes with a detection p-value >0.05 were excluded from further analysis.

Methylation data was analyzed using the SeSAMe package in R. Analysis was restricted to CpGs for which a gene annotation was available. Only CpGs that had β values for all subjects were retained.

A total of 347956 CpGs included a gene annotation in an age-associated module and were used in subsequent analysis. Correlation with age was determined using the “DML” function in SeSAMe, which takes a β value matrix and phenotype data as inputs.

To better distinguish genes whose expression is strongly under epigenetic control, we identified genes with a significant proportion of CpGs showing strong association with age. Fisher’s Exact Test was performed to find genes that were enriched with significant CpGs.

### Immunohistochemistry

FFPE samples were provided by the LTRC and were derived from subjects undergoing thoracic surgery. These subjects were diagnosed as being controls or having interstitial lung disease or COPD as determined by clinical history, CT scan, and surgical pathology. There was no intervention, as these are cross-sectional data. This dataset included 582 total subjects (254 have interstitial lung disease, 220 have COPD, and 108 are controls).

FFPE samples from healthy control human subjects were used for IHC. Samples were split into young and aged groups for immunohistochemistry, with 6 FFPE samples in each group. Age ranged from 44-53 years for young samples to 66-78 years for aged samples. FFPE blocks were processed as 5 micron thick sections.

**Table 1:** Subject demographic information for FFPE samples used for immunohistochemistry.

|  | Aged | Young |
| --- | --- | --- |
| Age (years) | 66 - 78 | 44 - 53 |
| Smoking History (>100 pkyr) | 3Y, 3N | 3Y, 3N |
| Sex | 3F, 3M | 3F, 3M |

Microwave antigen retrieval was performed using a pH 6.0 sodium citrate antigen retrieval buffer. SFTPC was detected using mouse anti-SFTPC polyclonal antibody diluted 1:200 (Santa Cruz Biotechnology, #518029) and HHIP was detected using rabbit anti-HHIP polyclonal antibody diluted 1:1500 (ABClonal Science, #A5872). Slides were incubated with primary antibody at 4°C for 16 hours, and later with HRP and AP Red secondary antibody for 1 hour. Immunoreactive signal was visualized with DAB solution (Vector Laboratories) and AP Red (Vector Laboratories). Slides were then counterstained with hematoxylin, dehydrated, and mounted.

To quantify staining, digital images of slides (40x magnification) were viewed using NIS-Elements (Nikon Instruments). Positive-stained cells were counted using color deconvolution in FIJI (National Institutes of Health). 5 fields per sample were obtained and cells were counted with a minimum size threshold of 300 pixels.

## Funding

The study was funded by NIH NHLBI grants R01HL127349, R01HL141852, U01HL145567, and UH2HL123886 to NK. RDM is an MD/PhD student with support from the Yale Medical Scientist Training Program (MSTP) grant (T32GM136651). EPM is a Pepper Scholar with support from the Claude D. Pepper Older Americans Independence Center at Yale School of Medicine (P30AG021342), Department of Veterans Affairs, Veterans Health Administration, VISN 1 Career Development Award, and NIA R03AG074063.

## Conflict of Interest

NK is a scientific founder at Thyron, served as a consultant to Biogen Idec, Boehringer Ingelheim, Third Rock, Pliant, Samumed, NuMedii, Theravance, LifeMax, Three Lake Partners, Optikira, Astra Zeneca, RohBar, Veracyte, Augmanity, CSL Behring, Galapagos, Fibrogen, and Thyron over the last 3 years, reports Equity in Pliant and Thyron, and grants from Veracyte, Boehringer Ingelheim, BMS and non-financial support from MiRagen and Astra Zeneca.

LN is the founder, President and CEO of Humacyte Global Inc, a publicly traded regenerative medicine company.

## Notes

https://www.ncbi.nlm.nih.gov/geo/query/acc.cgi?acc=GSE136831

https://www.ncbi.nlm.nih.gov/geo/query/acc.cgi?acc=GSE47460

https://gtexportal.org/home/datasets

## References

1 Bowdish, D. M. E. The Aging Lung Is Lung Health Good Health for Older Adults? Chest155, 391–400 (2019). https://doi.org:10.1016/j.chest.2018.09.003

2 Cho, S. J. & Stout-Delgado, H. W. Aging and Lung Disease. Annual Review of Physiology, Vol 82 82, 433–459 (2020). https://doi.org:10.1146/annurev-physiol-021119-034610

3 Chilosi, M., Poletti, V. & Rossi, A. The pathogenesis of COPD and IPF: Distinct horns of the same devil? Resp Res 13 (2012). https://doi.org:Artn 3 10.1186/1465-9921-13-3

4 Bartling, B. Cellular senescence in normal and premature lung aging. Z Gerontol Geriatr46, 613–622 (2013). https://doi.org:10.1007/s00391-013-0543-3

5 Budinger, G. R. S. et al. The Intersection of Aging Biology and the Pathobiology of Lung Diseases: A Joint NHLBI/NIA Workshop. J Gerontol A Biol Sci Med Sci 72, 1492–1500 (2017). https://doi.org:10.1093/gerona/glx090

6 Zeleznik, J. Normative aging of the respiratory system. Clin Geriatr Med 19, 1–18 (2003).

7 Copley, S. J. Morphology of the Aging Lung on Computed Tomography. J Thorac Imaging 31, 140–150 (2016). https://doi.org:10.1097/RTI.0000000000000211

8 Schroder, T. H., Storbeck, B., Rabe, K. F. & Weber, C. The Aging Lung: Clinical and Imaging Findings and the Fringe of Physiological State. Rofo 187, 430–439 (2015). https://doi.org:10.1055/s-0034-1399227

9 Brandenberger, C. & Muhlfeld, C. Mechanisms of lung aging. Cell Tissue Res 367, 469–480 (2017). https://doi.org:10.1007/s00441-016-2511-x

10 Bailey, K. L. et al. Aging causes a slowing in ciliary beat frequency, mediated by PKC epsilon. Am J Physiol-Lung C 306, L584–L589 (2014). https://doi.org:10.1152/ajplung.00175.2013

11 Godin, L. M. et al. Decreased Laminin Expression by Human Lung Epithelial Cells and Fibroblasts Cultured in Acellular Lung Scaffolds from Aged Mice. Plos One 11 (2016). https://doi.org:ARTN e0150966 10.1371/journal.pone.0150966

12 Lee, S. et al. Molecular programs of fibrotic change in aging human lung. Nat Commun 12 (2021). https://doi.org:ARTN 6309 10.1038/s41467-021-26603-2

13 Lonsdale, J. et al. The Genotype-Tissue Expression (GTEx) project. Nat Genet 45, 580–585 (2013). https://doi.org:10.1038/ng.2653

14 Saul, D. et al. A new gene set identifies senescent cells and predicts senescence-associated pathways across tissues. Nat Commun 13 (2022). https://doi.org:ARTN 4827 10.1038/s41467-022-32552-1

15 Xu, P. et al. The landscape of human tissue and cell type specific expression and coregulation of senescence genes. Mol Neurodegener 17 (2022). https://doi.org:ARTN 5 10.1186/s13024-021-00507-7

16 Chow, R. D., Majety, M. & Chen, S. D. The aging transcriptome and cellular landscape of the human lung in relation to SARS-CoV-2. Nat Commun 12 (2021). https://doi.org:ARTN 4 10.1038/s41467-020-20323-9

17 Angelidis, I. et al. An atlas of the aging lung mapped by single cell transcriptomics and deep tissue proteomics. Nat Commun 10 (2019). https://doi.org:ARTN 963 10.1038/s41467-019-08831-9

18 Ogrodnik, M., Salmonowicz, H. & Gladyshev, V. N. Integrating cellular senescence with the concept of damage accumulation in aging: Relevance for clearance of senescent cells (vol 18, e12841, 2019). Aging Cell 18 (2019). https://doi.org:ARTN e12942 10.1111/acel.12942

19 Langfelder, P. & Horvath, S. WGCNA: an R package for weighted correlation network analysis. Bmc Bioinformatics 9 (2008). https://doi.org:Artn 559 10.1186/1471-2105-9-559

20 Zhao, W. et al. Weighted Gene Coexpression Network Analysis: State of the Art. J Biopharm Stat 20, 281–300 (2010). https://doi.org:Pii 920109251 10.1080/10543400903572753

21 de Vries, M. et al. Lung tissue gene-expression signature for the ageing lung in COPD. Thorax 73, 609–617 (2018). https://doi.org:10.1136/thoraxjnl-2017-210074

22 Adams, T. S. et al. Single-cell RNA-seq reveals ectopic and aberrant lung-resident cell populations in idiopathic pulmonary fibrosis. Sci Adv 6 (2020). https://doi.org:ARTN eaba1983 10.1126/sciadv.aba1983

23 Hegab, A. E. et al. High fat diet activates adult mouse lung stem cells and accelerates several aging-induced effects. Stem Cell Res 33, 25–35 (2018). https://doi.org:10.1016/j.scr.2018.10.006

24 Lee, B. Y. et al. Senescence-associated beta-galactosidase is lysosomal beta-galactosidase. Aging Cell 5, 187–195 (2006). https://doi.org:10.1111/j.1474-9726.2006.00199.x

25 Peng, Q., Gao, L., Cheng, H. B., Wang, J. S. & Wang, J. Sialidase NEU1 May Serve as a Potential Biomarker of Proliferation, Migration and Prognosis in Melanoma. World J Oncol 13, 222–234 (2022). https://doi.org:10.14740/wjon1509

26 Sauler, M. et al. Characterization of the COPD alveolar niche using single-cell RNA sequencing. Nat Commun 13 (2022). https://doi.org:ARTN 494 10.1038/s41467-022-28062-9

27 McDonough, J. E. et al. Low Surfactant Type II Alveolar Epithelial Cells Are an Enriched Cell Population in the Aged Lung. Am J Resp Crit Care 201 (2020).

28 Travaglini, K. J. et al. A molecular cell atlas of the human lung from single-cell RNA sequencing. Nature 587 (2020). https://doi.org:10.1038/s41586-020-2922-4

29 Avelar, R. A. et al. A multidimensional systems biology analysis of cellular senescence in aging and disease. Genome Biol 21 (2020). https://doi.org:ARTN 91 10.1186/s13059-020-01990-9

30 Hernandez-Segura, A. et al. Unmasking Transcriptional Heterogeneity in Senescent Cells. Curr Biol 27, 2652−+ (2017). https://doi.org:10.1016/j.cub.2017.07.033

31 Negrini, S., Gorgoulis, V. G. & Halazonetis, T. D. Genomic instability - an evolving hallmark of cancer. Nat Rev Mol Cell Bio 11, 220–228 (2010). https://doi.org:10.1038/nrm2858

32 Garcia-Nieto, P. E., Morrison, A. J. & Fraser, H. B. The somatic mutation landscape of the human body. Genome Biol 20 (2019). https://doi.org:ARTN 298 10.1186/s13059-019-1919-5

33 Shi, J. J., Surma, M., Yang, Y. & Wei, L. Disruption of both ROCK1 and ROCK2 genes in cardiomyocytes promotes autophagy and reduces cardiac fibrosis during aging. Faseb J 33, 7348–7362 (2019). https://doi.org:10.1096/fj.201802510R

34 Zhang, Y. J. et al. TIPARP is involved in the regulation of intraocular pressure. Commun Biol 5 (2022). https://doi.org:ARTN 1386 10.1038/s42003-022-04346-0

35 Huang, W. J., Hickson, L. J., Eirin, A., Kirkland, J. L. & Lerman, L. O. Cellular senescence: the good, the bad and the unknown. Nat Rev Nephrol 18, 611–627 (2022). https://doi.org:10.1038/s41581-022-00601-z

36 Lao, T. et al. Haploinsufficiency of Hedgehog interacting protein causes increased emphysema induced by cigarette smoke through network rewiring. Genome Med 7, 12 (2015). https://doi.org:10.1186/s13073-015-0137-3

37 Zhou, X. et al. Identification of a chronic obstructive pulmonary disease genetic determinant that regulates HHIP. Hum Mol Genet 21, 1325–1335 (2012). https://doi.org:10.1093/hmg/ddr569

38 Lao, T. et al. Hhip haploinsufficiency sensitizes mice to age-related emphysema. Proc Natl Acad Sci U S A 113, E4681–4687 (2016). https://doi.org:10.1073/pnas.1602342113

39 Schousboe, P. et al. Reduced levels of pulmonary surfactant in COVID-19 ARDS. Sci Rep-Uk 12 (2022). https://doi.org:ARTN 4040 10.1038/s41598-022-07944-4

40 Lopez-Otin, C., Blasco, M. A., Partridge, L., Serrano, M. & Kroemer, G. The Hallmarks of Aging. Cell 153, 1194–1217 (2013). https://doi.org:10.1016/j.cell.2013.05.039

41 Yang, J. L. et al. Synchronized age-related gene expression changes across multiple tissues in human and the link to complex diseases. Sci Rep-Uk 5 (2015). https://doi.org:ARTN 15145 10.1038/srep15145

42 Kumari, R. & Jat, P. Mechanisms of Cellular Senescence: Cell Cycle Arrest and Senescence Associated Secretory Phenotype. Front Cell Dev Biol 09 (2021). https://doi.org:ARTN 645593 10.3389/fcell.2021.645593

43 Ngassie, M. L. K. et al. Age-associated Differences in the Human Lung Extracellular Matrix. bioRxiv, 2022.2006.2016.496465 (2023).https://doi.org:10.1101/2022.06.16.496465

44 Tuttle, C. S. L. et al. Cellular senescence and chronological age in various human tissues: A systematic review and meta-analysis. Aging Cell 19 (2020). https://doi.org:ARTN e13083 10.1111/acel.13083

45 Chen, J. H., Hales, C. N. & Ozanne, S. E. DNA damage, cellular senescence and organismal ageing: causal or correlative? Nucleic Acids Res 35, 7417–7428 (2007). https://doi.org:10.1093/nar/gkm681

46 von Zglinicki, T., Wan, T. & Miwa, S. Senescence in Post-Mitotic Cells: A Driver of Aging? Antioxid Redox Signal 34, 308–323 (2021). https://doi.org:10.1089/ars.2020.8048

47 Sozou, P. D. & Kirkwood, T. B. L. A stochastic model of cell replicative senescence based on telomere shortening, oxidative stress, and somatic mutations in nuclear and mitochondrial DNA. J Theor Biol 213, 573–586 (2001). https://doi.org:10.1006/jtbi.2001.2432

48 Tsuji, T., Aoshiba, K. & Nagai, A. Alveolar cell senescence in patients with pulmonary emphysema. Am J Resp Crit Care 174, 886–893 (2006). https://doi.org:10.1164/rccm.200509-1374OC

49 Enge, M. et al. Single-Cell Analysis of Human Pancreas Reveals Transcriptional Signatures of Aging and Somatic Mutation Patterns. Cell 171, 321−+ (2017). https://doi.org:10.1016/j.cell.2017.09.004

50 Crouch, J., Shvedova, M., Thanapaul, R. J. R. S., Botchkarev, V. & Roh, D. Epigenetic Regulation of Cellular Senescence. Cells-Basel 11 (2022). https://doi.org:ARTN 672 10.3390/cells11040672

51 Wang, X. R., Park, J., Susztak, K., Zhang, N. R. & Li, M. Y. Bulk tissue cell type deconvolution with multi-subject single-cell expression reference. Nat Commun 10 (2019). https://doi.org:ARTN 380 10.1038/s41467-018-08023-x

52 McDonough, J. E. et al. Transcriptional regulatory model of fibrosis progression in the human lung. JCI Insight 4 (2019). https://doi.org:10.1172/jci.insight.131597

53 McDonough, J. E. et al. Small-airway obstruction and emphysema in chronic obstructive pulmonary disease. N Engl J Med 365, 1567–1575 (2011). https://doi.org:10.1056/NEJMoa1106955

